# Optineurin Deficiency Collapses the Host Endolysosomal Network and Impairs Xenophagy to Accelerate *Mycobacterium tuberculosis* Growth

**DOI:** 10.64898/2026.08.23.746595

**Authors:** Nalin Abeydeera, Jeffrey Chin, Angel Ruvalcaba, Khushi Amin, Vinh Q Nguyen, Joshua Chang, Katherine Yin, Joel D Ernst, Jonathan M Budzik

## Abstract

Selective autophagy is a host defense mechanism against *Mycobacterium tuberculosis* (*Mtb*) that restricts bacterial growth by targeting ubiquitin-coated bacilli for lysosomal degradation via autophagy receptors. Optineurin is a selective autophagy receptor that targets pathogens and modulates immune signaling; however, its precise structural mechanism during *Mtb* infection remains poorly defined. Here, we show that while Optineurin deficiency spares the global host transcriptomic response to infection, it collapses the host endolysosomal network, reducing LAMP1⁺ and LysoTracker⁺ reserves by half and 30%, respectively. Multi-dose bafilomycin A1 flux assays demonstrated that this structural depletion selectively blocks the dynamic, directional trafficking and functional delivery of autophagosomes to the pathogen, significantly reducing *Mtb*-DQ-BSA colocalization. Genetic complementation restored bacterial restriction in a manner dependent on three phosphosites (Ser187, Ser530, and the uncharacterized Ser556). In the context of reduced autophagic containment and increased *Mtb* replication, Optineurin deficiency accelerated necrotic-like host cell death. *In vivo*, Optineurin deficiency enhanced bacterial replication and impaired the Type I interferon response during acute *Mtb* infection but did not affect long-term survival. Together, these findings identify Optineurin as a critical regulator of autophagic flux, host cell death, and Type I interferon responses that limit early *Mtb* pathogenesis.

**Importance:** Tuberculosis remains a major global health threat in part because *Mycobacterium tuberculosis* can survive and replicate inside immune cells meant to destroy it. Host cells use a specialized cellular recycling and defense system, called autophagy, to capture and eliminate these intracellular bacteria. In this study, we identified a critical cellular protein, Optineurin, that serves as an essential sensor initiating this defense mechanism during infection. We discovered that chemical modifications, specifically phosphorylation, act as molecular switches that activate Optineurin’s protective functions. Without this protein, host cells control bacterial growth less effectively, suffer premature cell death, and partially lose their ability to mount early cytokine responses. By uncovering how Optineurin coordinates these diverse defense pathways, our findings provide a deeper understanding of early host-pathogen interactions and reveal potential molecular targets for developing novel, host-directed therapies to combat tuberculosis.

## Introduction

*Mycobacterium tuberculosis* (*Mtb*) remains a global health threat, as there is no vaccine that provides consistent, long-term adult protection (1). Upon inhalation, *Mtb* primarily infects and replicates within alveolar macrophages (2, 3). Subsequently, the bacilli traffic into recruited monocyte-derived macrophages and neutrophils (4). Despite the antimicrobial effects of adaptive interferon-gamma (IFN-y), *Mtb* utilizes sophisticated counter-mechanisms to survive and replicate within these host cell subsets (5–9).

Xenophagy is a primary cell-intrinsic host defense strategy that *Mtb* actively circumvents. In this selective macroautophagy pathway, intracellular pathogens are decorated with lysine-48 (K48) and lysine-63 (K63) polyubiquitin chains (10, 11). Specialized autophagy receptors recognize and bridge the ubiquitinated cargo to recruit autophagosomal machinery via microtubule-associated protein 1A/1B-light chain 3 (LC3) (12). Following lipidation, lipid-conjugated LC3-II drives autophagosome membrane formation around the pathogen (13), culminating in fusion with lysosomes to generate degradative autolysosomes (14–19). Highlighting its evolutionary importance, *Mtb* deploys virulence factors that inhibit autophagic maturation (20), halt phagosome-lysosome fusion (21–23), and arrest luminal acidification (24–27).

The physiological requirement for this pathway is underscored by the fact that macrophages deficient in core autophagy-related genes (*Atg5*, *Atg16L1*, *Atg7)* display significantly enhanced *Mtb* replication (28, 29) and undergo an accelerated kinetic shift toward necrotic-like host cell death (29), which promotes bacterial dissemination (29–32). Systemic Atg5 deficiency dramatically exacerbates *Mtb* pathogenesis (17, 29, 33, 34), whereas Atg16L and Atg7 deficiencies exhibit more moderate phenotypes (29, 33). Conversely, deficiency of another gene that mediates autophagy, Tax1bp1, restricts *Mtb* growth *in vivo,* revealing that autophagy receptor-mediated regulation of collateral immune responses, beyond degradative clearance, shapes pathogenesis (16, 35).

Over 30 selective autophagy receptors exist, each tailored to specific intracellular cargoes (36). Four are known to recognize ubiquitinated *Mtb*: Ndp52 (17), p62/SQSTM1 (17, 37), Tax1bp1 (14–16), and Optineurin (14, 15, 19). To orchestrate cargo clearance, these receptors undergo post-translational modifications that tune their activation state. Notably, site-specific phosphorylation of core functional domains enhances their affinity for LC3 and polyubiquitin (38–47).

Optineurin is a versatile autophagy receptor with established roles in innate immunity. Following recruitment to ubiquitinated *Salmonella enterica,* phosphorylation of Optineurin at Ser177 by TANK-binding kinase 1 (TBK1) enhances its LC3-binding affinity (40). Consequently, cells expressing point-mutants that abrogate ubiquitin or LC3 binding fail to restrict *Salmonella* (40). Similarly, Optineurin deficiency elevates the bacterial burden during *Mycobacterium marinum* infection and reduces LC3-pathogen colocalization (48). Optineurin depletion also accelerates *Listeria monocytogenes* replication (49, 50). Finally, baseline LC3-II conversion is decreased in Optineurin-deficient BMDMs (bone marrow-derived macrophages) infected with *Mycobacterium smegmatis* (51), establishing a protective role for Optineurin across various bacterial species.

Beyond xenophagy, Optineurin clears diverse endogenous cargoes. By binding to Rab8 and Myosin VI, Optineurin acts as a mechanical tether to drive the autophagic clearance of neurodegenerative protein aggregates (52–54), functionally linking its disruption to the pathogenesis and chronic progression of Amyotrophic Lateral Sclerosis and glaucoma (55). Optineurin also coordinates organelle homeostasis by activating transcription factor EB (TFEB) to drive baseline lysosomal biogenesis in the retinal pigment epithelium (56) and by regulating mitophagy and starvation-induced non-selective autophagy (57–59). A growing body of literature describes highly cell-type– and stimulus-dependent specificity in the effects of Optineurin deficiency on starvation dynamics, showing either diminished autophagosome formation or an accumulation of vesicles resulting from a block in lysosomal fusion (57–59). Conversely, mitophagy-induced HeLa cells maintain stable LC3-II levels despite Optineurin deficiency (60), revealing variable effects on autophagosome formation versus downstream clearance kinetics.

We and others have shown that Optineurin colocalizes with the *Mtb* vacuole (14, 15, 19) and is phosphorylated during infection (15). Because cargo encapsulation and phosphorylation denote receptor activation (61), we hypothesized that Optineurin regulates virulent *Mtb* autophagic flux and that site-specific domain phosphorylation is required for host restriction. Using genetic complementation, we demonstrate that Optineurin deficiency accelerates virulent *Mtb* growth in macrophages, a phenotype reversed by transgenic restoration of an *Optineurin* complementation allele. Strikingly, while Optineurin deficiency spares global host transcriptomic responses and the initial upstream recruitment of polyubiquitin and collateral receptors (Tax1bp1, p62) to the bacilli, it collapses the host endolysosomal network, decreasing structural LAMP1+ and acidified LysoTracker+ reserves by half and 30%, respectively. Multi-dose bafilomycin A1 flux assays reveal that this depletion selectively blocks the downstream, directional targeting of autophagosomes to the pathogen vacuole, culminating in accelerated macrophage necrosis. Furthermore, alanine mutagenesis identified three independent phosphosites, including the previously uncharacterized Ser556 residue in the zinc finger domain, individually required for bacterial restriction. Finally, *in vivo* aerosol infection of Optineurin-deficient mice recapitulated the enhanced pathogen growth phenotype, driving expansion of pulmonary *Mtb* burden, systemic dissemination, and an altered IL-1β^hi^/IFN-β^lo^ immune microenvironment during acute infection without shortening survival. Together, our study provides the first demonstration that Optineurin restricts virulent *Mtb* pathogenesis in macrophages and *in vivo* via a mechanism governed by multi-domain phosphorylation.

## Results

### Optineurin restricts the growth of *Mtb* in macrophages

To evaluate Optineurin function during *Mtb* infection, we engineered an Optineurin-deficient conditionally immortalized macrophage (CIM) line (62). Cas9-expressing CIMs were transduced with a single-guide RNA (sgRNA) targeting exon 2 of *Optineurin*, and single-cell limiting dilution established a monoclonal cell line with 100% editing efficiency.

CRISPR editing was validated via PCR, Sanger sequencing, and Inference of CRISPR edits (ICE) analysis (63), revealing a single-nucleotide (–1) cytosine deletion at position 109 of the coding sequence (Fig. S1). This mutation introduces a premature stop codon after 43 amino acids, targeting the transcript for nonsense-mediated decay. To control for off-target effects (64), the knockout line was complemented via lentiviral transduction with an *Optineurin* allele harboring synonymous mutations in the sgRNA-binding sequence and protospacer adjacent motif (PAM) to shield the transgene from Cas9 cleavage (Fig. 1A).

**Fig. 1.**
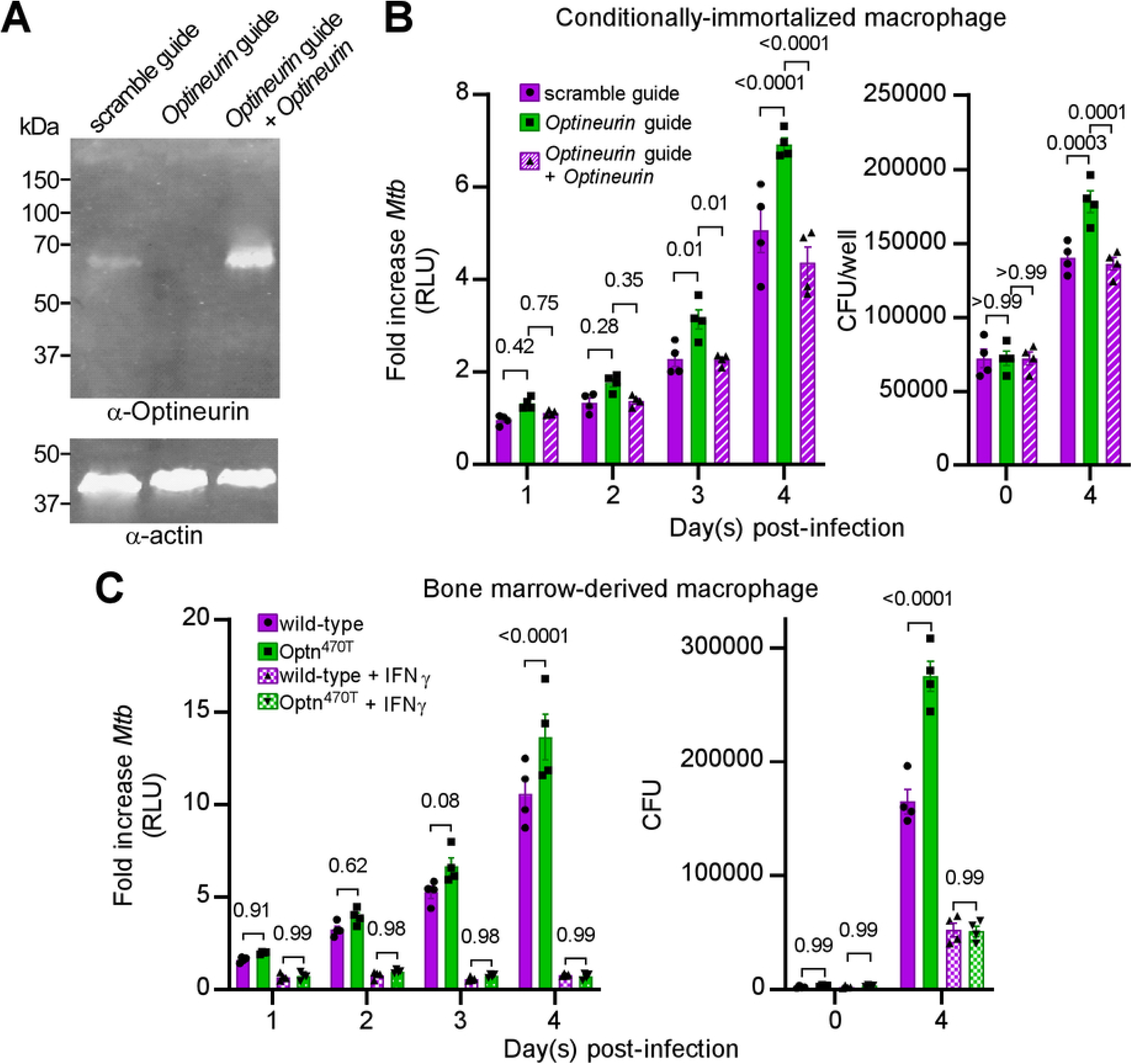
Optineurin restricts *Mtb* growth in macrophages. (A) CIMs were transduced with lentivirus expressing sgRNA targeting *Optineurin* (or scramble control sgRNA) without or with Flag-Optineurin. Cellular lysates were subjected to SDS-PAGE (Sodium dodecyl sulfate polyacrylamide gel electrophoresis) and immunoblotted with primary antibodies against Optineurin or actin and infrared-dye-conjugated secondary antibodies. Fluorescence images are presented. The relative mobilities of the molecular weight markers are displayed. (B) CIMs were infected with luciferase-expressing *Mtb* H37Rv (MOI 1). In four wells per condition, the infected monolayers were lysed immediately after infection and plated for colony-forming units (CFU) (day 0). In the remaining wells, the cell culture media were replaced daily with fresh media. Monolayer luminescence was measured daily. Following lysis of the monolayers, CFU were measured four days after infection. (C) BMDMs were infected with *Mtb* Erdman (MOI 1). The media were replaced daily with fresh media with or without IFN-y. Monolayer luminescence was measured daily. CFU were measured 4 days after infection. The mean, SEM, and adjusted p-values from two-way ANOVA (Tukey’s multiple comparison test) are displayed. Data are representative of two independent experiments.

Infected macrophage lines were tracked daily to measure intracellular replication. Optineurin deficiency accelerated *Mtb* replication, yielding a 37% greater fold-increase in monolayer bioluminescence (Fig. 1B, left) and 1.27-fold increase in colony-forming units (CFUs) at 4 days post-infection (Fig. 1B, right). Transgenic expression of the complemented *Optineurin* allele effectively reversed this phenotype, restoring bacterial restriction to levels observed in the scramble control.

To validate these phenotypes in primary cells, we quantified infection kinetics in Optn^470T^ BMDMs expressing a mutant Optineurin lacking the C-terminal UBAN and zinc finger regions (Fig. 1C) (65). Compared to wild-type controls, infected Optn^470T^ BMDMs displayed accelerated replication, with a 29% greater fold-increase in bioluminescence and 1.67-fold increase in CFUs at 4 days post-infection (Fig. 1C). Crucially, Optineurin-deficient BMDMs remained fully capable of restricting *Mtb* growth when activated with exogenous interferon-gamma (IFN-y; Fig. 1C). Thus, Optineurin is a cell-intrinsic restriction factor that controls *Mtb* replication.

### Optineurin deficiency reduces autophagy flux of *Mtb*

Given that Optineurin is a selective autophagy receptor (40), we tested whether enhanced bacterial growth was driven by reduced pathogen degradation via selective macroautophagy (xenophagy). We quantified the physical overlap of fluorescent *Mtb* with essential components of the ubiquitin-dependent xenophagy cascade in wild-type and Optn^470T^ BMDMs (Fig. 2, Fig. S2). At 8 and 24 h post-infection, we observed no statistically significant differences in the colocalization of *Mtb* with phosphorylated TBK1 (p-TBK1), polyubiquitin, p62, Tax1bp1, or LC3 (Fig. 2E-F, Fig. S2). This invariant baseline indicates that Optineurin deficiency does not compromise upstream pathogen surveillance, cargo recognition, or autophagosome initiation at steady state.

**Fig. 2.**
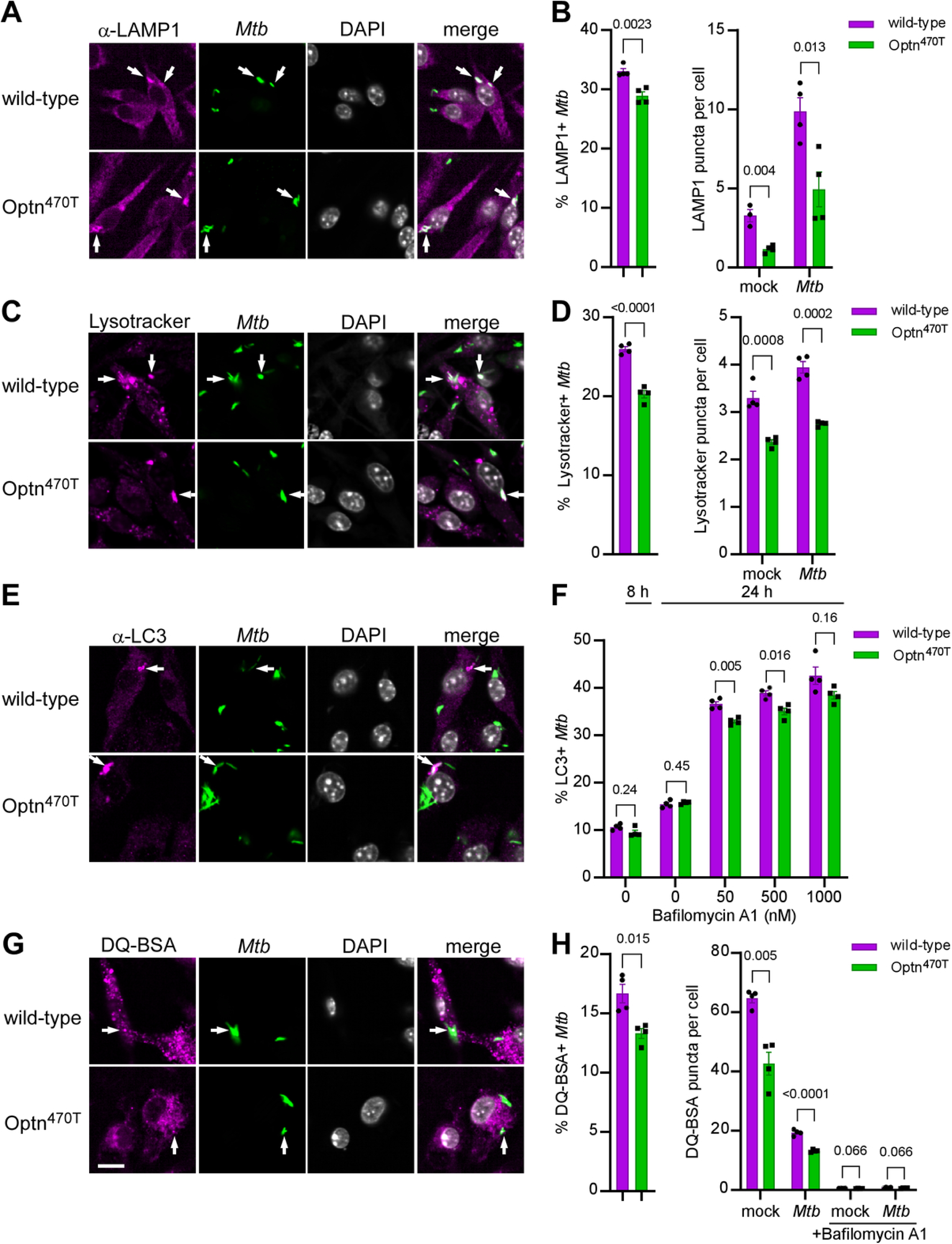
Optineurin deficiency reduces autophagy flux in *Mtb*. BMDMs were infected with ZsGreen-expressing *Mtb* (MOI 2) or mock-infected. (A-B) Macrophages were fixed at 8 h post-infection and stained with primary antibodies for LAMP1, secondary Alexa-Fluor 647 antibodies, and DAPI. (C-D) Macrophages were treated with Lysotracker at 5 h post-infection, and the monolayers were fixed at 8 h post-infection, stained with DAPI, and immediately imaged using confocal microscopy. (E-F) *Mtb-*infected BMDMs were fixed at 8 h post-infection or separately treated with bafilomycin for 3 h prior to fixation at 24 h post-infection. The samples were stained with primary antibodies for LC3, secondary Alexa-Fluor 647 antibodies, and DAPI. A representative image from 24 h post-infection (0 nM bafilomycin) is displayed in panel (E). (G-H) Macrophages were treated with bafilomycin at 5 h post-infection, DQ-BSA was added at 7 h post-infection, and the monolayers were fixed at 8 h post-infection. Immunofluorescence microscopy was performed at 63 × magnification in a minimum of 86 x/y positions and four z planes each in quadruplicate wells per experimental condition. At least 2000 macrophages were imaged per well. Arrows in the representative images (A, C, E, G) denote *Mtb* colocalization with antibody or probe puncta. The white bar denotes 10 µm. (B, D, F, H) Quantification of colocalization with *Mtb* and antibody or probe puncta, and the number of puncta (LAMP1, Lysotracker, or DQ-BSA) are shown. (B, D, H) Mean percent colocalization in each well, SEM, and p-values from the unpaired, two-tailed t-test are displayed. Mean percent colocalization in each well (F) or mean fluorescent puncta per cell in each well (B, D, H), SEM, and p-values from the unpaired, two-tailed t-tests with Holm-Sidak multiple comparison correction are displayed. Data are representative of two (Fig. 2A-F) or four (Fig. 2G-H) independent experiments.

However, because Optineurin mediates lysosomal quality control (66), and *Mtb* inflicts structural damage on endolysosomal membranes (67), we reasoned that a steady-state snapshot might mask a kinetic bottleneck. Moreover, as lysosomal quality and repair are prerequisites for sustained autophagic flux (68, 69), we hypothesized that Optineurin deficiency compromises the dynamic flux of *Mtb* autophagic processing, impairing downstream delivery and clearance.

To test this, we independently assessed lysosomal abundance, acidification, and proteolytic activity. We first quantified total lysosomal membrane infrastructure using immunofluorescence staining for the static membrane marker LAMP1 at 8 h post-infection. High-content imaging analysis revealed that Optineurin deficiency compromises lysosomal abundance under both uninfected and infected conditions. In uninfected cells, the mean number of LAMP1 puncta per cell dropped from 3.28 in wild-type controls to 1.15 in mutant cells (Fig. 2A-B; Fig. S3A). Under infection stress, this deficit persisted, whereby LAMP1 abundance decreased by half from a mean of 9.86 puncta per cell in wild-type cells to 4.94 puncta per cell in mutants (Fig. 2A-B). This pool depletion was accompanied by a decrease in physical pathogen targeting, with *Mtb-* LAMP1 colocalization shifting from 32% in wild-type cells to 28% in mutant cells (Fig. 2A-B).

To determine whether this loss of infrastructure caused an acidification defect, we quantified the total number of acidified vesicles using the acidotropic dye LysoTracker at 8 h post-infection. In the uninfected condition, the total LysoTracker puncta per cell decreased from a mean of 3.30 in wild-type cells to 2.37 in mutants (Fig. 2C-D; Fig. S3A). Following *Mtb* infection, a comparable 29.5% decrease occurred, decreasing from a mean of 3.90 puncta per cell in wild-type macrophages to 2.75 in mutants (Fig. 2C-D). The colocalization of *Mtb* and LysoTracker also decreased from 26% in wild-type cells to 21% in mutant cells (Fig. 2C-D).

To map how the depleted endolysosomal reserve intersects with host defense pathways, we evaluated autophagosome formation and pathogen targeting via LC3 imaging at 24 h post-infection across a range of bafilomycin A1 concentrations. At baseline without a chemical clamp, mutant cells exhibited slightly lower bulk autophagosome protein density compared to wild-type cells, with mean corrected LC3 spot intensity decreasing from 40.9 to 38.4 (Fig. S3B-C). Interestingly, baseline physical autophagosome abundance was identical between genotypes, with both wild-type and mutant macrophages displaying a mean of 2.1 LC3 puncta per cell (Fig. S3B-C), while the baseline LC3 percent colocalization with *Mtb* remained equivalent (10.6% in wild-type versus 9.5% in Optineurin-deficient cells at 8 h, and 15.4% in wild-type versus 15.8% in Optineurin-deficient cells at 24 h; Fig. 2E-F).

To resolve dynamic autophagic flux versus vesicle biogenesis, cells were treated with the V-ATPase inhibitor bafilomycin A1, which blocks the degradation of LC3-II in acidic lysosomes (70). Across all tested doses, the number of LC3 puncta per cell rose uniformly and showed no statistically significant differences between genotypes, shifting from 6.7 versus 6.0 puncta at 50 nM, 7.9 versus 7.2 puncta at 500 nM, and 8.8 versus 8.5 puncta at 1000 nM in wild-type and mutant cells, respectively (Fig. S3B-C). This uniform puncta accumulation demonstrates that Optineurin deficiency does not impair upstream autophagosome biogenesis.

Crucially, while autophagosomes remained constant across genotypes, spatial analysis revealed a significant, selective defect in pathogen targeting. Under a 50 nM clamp, despite an equivalent number of vesicles, mutant macrophages displayed decreased targeted pathogen capture compared to wild-type controls, with LC3-*Mtb* colocalization decreasing from 36.6% to 32.9% (Fig. 2E-F; Fig. S3B). This selective targeting defect was mirrored at 500 nM, where colocalization dropped from 38.9% in wild-type cells to 35.1% in mutants (Fig. 2E-F; Fig S3B). Corrected spot intensities tracked with these shifts (Fig. S3C), with wild-type values at a mean of 69.0 versus 60.0 at 50 nM and 78.0 versus 67.7 at 500 nM. A saturating 1000 nM bafilomycin A1 block rendered bulk intensity differences non-significant (93.4 in wild-type vs. 77.6 in mutant) while preserving the downward trend in pathogen targeting (42.6% in wild-type vs. 38.5% in mutant). Collectively, these multi-dose flux assays demonstrate that Optineurin deficiency compromises autophagosomal cargo clearance and directional targeting to the pathogen vacuole rather than disrupting baseline autophagosome formation.

Thus, while Optineurin deficiency depletes the permanent host lysosomal reserve (LAMP1 and LysoTracker puncta), equivalent LC3 puncta across all conditions demonstrates that upstream autophagosome biogenesis remains intact. The host defect is characterized by the collapse of the lysosomal structural network and the inability to direct these normally formed autophagosomes to the pathogen vacuole.

### Optineurin deficiency impairs proteolytic capacity

To determine whether the structural depletion of the host endolysosomal network led to a loss of cargo processing, we assessed lysosomal enzymatic cleavage using the fluorogenic protease substrate DQ-BSA. Macrophages were pulsed with DQ-BSA at 7 h post-infection and fixed at 8 h post-infection to capture protease-dependent fluorescence unquenching. High-content imaging demonstrated that Optineurin deficiency compromises baseline proteolytic capacity. Under uninfected (mock) conditions, Optn^470T^ macrophages exhibited a significant reduction in global lysosomal activity, with DQ-BSA corrected spot intensity decreasing from a mean of 303.1 in wild-type cells to 270.8 in mutants (Fig. S4A,C). This global enzymatic deficit was mirrored by a reduction in total active vesicle abundance under mock baseline conditions, where the mean number of DQ-BSA puncta per cell decreased from 64.7 in wild-type cells to 42.6 in mutants (Fig. 2H).

Intriguingly, *Mtb* infection imposed a dominant layer of metabolic suppression that overrode these baseline genetic differences. Upon infection, global DQ-BSA-corrected spot intensities decreased to a shared baseline in both wild-type and mutant cells (means of 156.6 vs. 157.2, respectively; Fig. S4B-C), consistent with *Mtb*-mediated V-ATPase exclusion and intraphagosomal deacidification. A parallel drop occurred in total active vesicle numbers, decreasing to 19.3 puncta per cell in infected wild-type cells and to 13.3 in mutants (Fig. 2H, Fig. S4B).

Next, we evaluated how this infrastructure collapse and global enzymatic suppression altered the functional delivery of active proteases to the pathogen vacuole. In wild-type macrophages, baseline colocalization between *Mtb* and active DQ-BSA compartments was expectedly constrained at a mean of 16.7%, reflecting *Mtb*’s intrinsic capacity to arrest phagosome maturation (Fig. 2G-H). Remarkably, Optineurin deficiency drove a further reduction in functional cargo delivery, decreasing *Mtb-*DQ-BSA colocalization to a mean of 13.3% (Fig. 2G-H). This significant functional targeting defect mirrors the downward shifts observed independently for structural LAMP1 membranes (Fig. 2A-B) and acidified LysoTracker compartments (Fig. 2C-D).

To validate the probe’s acidification dependence and evaluate vesicle dynamics under a global fusion block, controls were treated with bafilomycin A1 at 5 h post-infection. In uninfected samples, bafilomycin collapsed total active puncta to near-zero levels that were indistinguishable between genotypes (0.53 in wild-type vs. 0.41 in mutant; Fig. 2H, Fig. S4B) and equalized global corrected spot intensities (means of 134.0 vs. 145.6, respectively; Fig. S4A,C). This bafilomycin-mediated suppression remained uniformly robust under infected conditions, yielding statistically equivalent residual puncta counts (0.75 vs 0.54 puncta/cell; Fig. 2H, Fig. S4B) and corrected spot intensities (means of 165.5 vs 174.5; Fig. S4B-C) between genotypes. This uniform drug action confirms that Optineurin deficiency does not alter cellular sensitivity to V-ATPase inhibition, thereby validating that the observed baseline defects represent a specific, cell-intrinsic collapse of the functional endolysosomal reserve.

### Optineurin deficiency does not impact bulk metabolic autophagy in BMDMs

Because Optineurin deficiency reduces *Mtb* autophagic flux, we investigated whether it plays a broader role in non-selective bulk macroautophagy in macrophages. Wild-type and Optineurin-deficient BMDMs were subjected to nutrient starvation via Earle’s Balanced Salt Solution (EBSS) alongside bafilomycin A1 treatment to assess dynamic autophagic flux (71–74). Immunoblot analysis revealed a robust increase in the LC3-II/LC3-I ratio under nutrient-limited conditions (Fig. S5A). However, no statistically significant differences in the LC3-II/LC3-I conversion ratio were observed between genotypes under basal or starvation and bafilomycin conditions (Fig. S5B). These bulk biochemical findings demonstrate that Optineurin is dispensable for non-selective bulk macroautophagy, indicating its role is specialized for pathogen-directed xenophagy.

### The bulk transcriptional responses to *Mtb* are similar in wild-type and Optineurin-deficient macrophages

Because Optineurin deficiency has been linked to disrupted TFEB-mediated transcription (56), we used global RNA sequencing (RNA-seq) on mock– or *Mtb*-infected wild-type and Optn^470T^ BMDMs across three independent replicates to test whether our observed lysosomal deficits were transcriptionally driven.

Principal component analysis (PCA) confirmed that pathogen exposure was the primary driver of transcriptional variance, with PC1 explaining 44.6% of the variance and segregating mock-infected from *Mtb*-infected samples (Fig. S6A). PC2 accounted for 20.1% of the variance, reflecting the expected variation across independent primary BMDM harvests (Fig. S6A). Crucially, the wild-type and mutant genotypes within each independent biological replicate were paired and clustered (Fig. S6A), indicating no independent transcriptional divergence between genotypes.

Differential gene expression analysis revealed that *Mtb* infection triggered extensive, similar transcriptional reprogramming in both genotypes. Examples of upregulated transcripts with high fold-change that were significantly enriched included genes driving antigen presentation (*H2-M2*) (75, 76), inflammation (C*xcl1*, *Cxcl3*, *Cxcl10*, *Cxcl11*) (77, 78), leukocyte chemotaxis (*Fpr1*) (79), pro-inflammatory cytokine responses (*Il6*, *Tarm1*) (80), tissue homeostasis (*Orm1*, *Lox*) (81, 82), and Type I interferon responses (*Ifnb1*) (83) (Fig. S6B-C, yellow puncta). Conversely, transcripts enriched in mock-infected samples included a pattern recognition receptor (*Mrc1*) (84, 85) as well as a regulator of endophagosomal acidification (*Slc9a9*) (86) (Fig. S6B-C, blue puncta).

Venn diagram analysis confirmed substantial transcriptional conservation, with 245 upregulated (Fig. S6F) and 50 downregulated (Fig. S6G) genes shared symmetrically between genotypes. Direct pairwise comparisons revealed zero differentially expressed genes between wild-type and Optineurin-deficient macrophages under mock-infected (Fig. S6E) or *Mtb*-infected (Fig. S6D) conditions. These analyses demonstrate that Optineurin deficiency does not alter global mRNA abundance or compromise transcriptional induction under these experimental conditions, confirming that depletion of the endolysosomal network and autophagic trafficking bottlenecks operate at the post-transcriptional level. It remains open, however, whether Optineurin deficiency selectively drives transcriptional shifts within distinct BMDM subpopulations, such as highly infected versus bystander cells, a nuance that requires single-cell RNA sequencing to fully uncover (87–90).

### Expression of phosphosite-deficient Optineurin enhances *Mtb* growth

As one example of a mechanism operating at the post-transcriptional level, we focused on post-translational modifications. We previously discovered that Optineurin is phosphorylated at multiple sites during *Mtb* infection: S187 in the LC3-interacting region (LIR), a cluster at T210/T212/T214, S530 within the ubiquitin-binding region (UBAN), and S556 within the zinc finger domain (15). Given their positions within key functional domains, we hypothesized that these site-specific phosphorylation events mediate *Mtb* restriction. We performed growth assays in *Optineurin-*edited CIMs engineered to express non-phosphorylatable alanine substitution alleles, with successful expression validated by immunoblotting (Fig. S7).

Expression of single phosphosite-deficient alleles at residues S187A, S530A, or S556A significantly accelerated *Mtb* growth, as measured by both bioluminescence (Fig. 3A) and CFU assays (Fig. 3B). In contrast, the triple phosphomutant allele (Optineurin^T210A/T212A/T214A^) matched the restriction capacity of controls transduced with the scramble guide (Fig. 3A-B). Together, these results demonstrate that phosphorylation at three distinct functional domains (the LIR motif, the UBAN domain, and the zinc finger domain) is individually required for Optineurin-mediated restriction of *Mtb*.

**Fig. 3.**
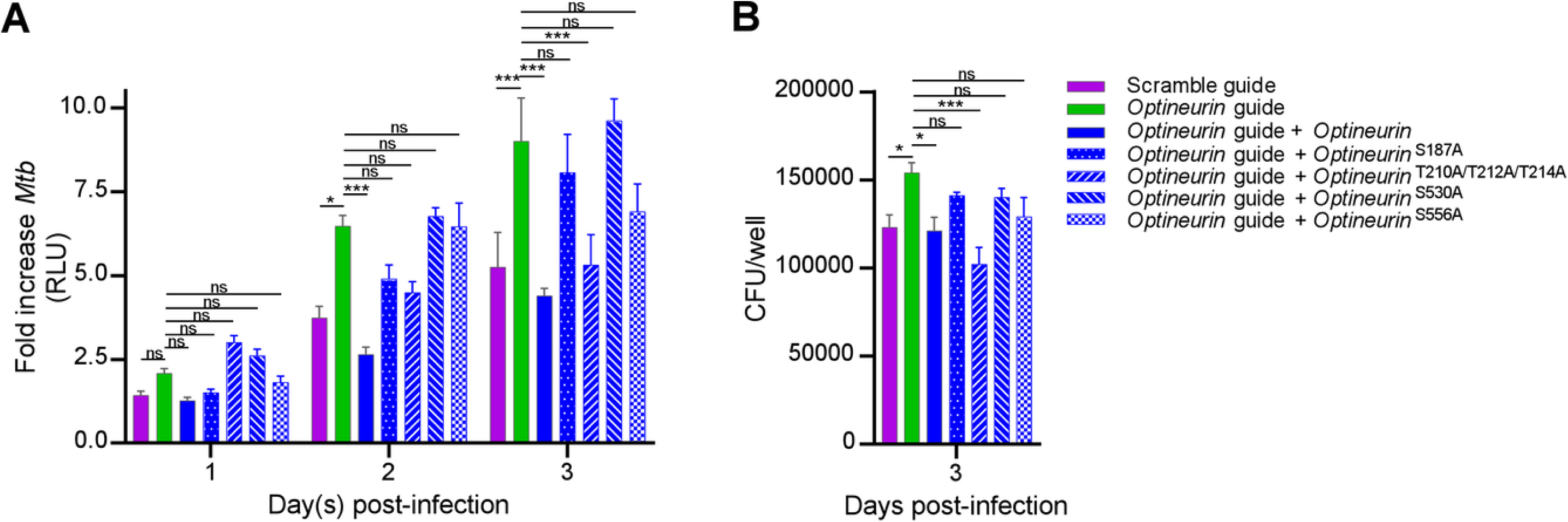
Phosphosite-deficient Optineurin at Ser187, Ser513, and Ser556 restrict *Mtb* growth. CIMs transduced with lentivirus expressing sgRNA targeting *Optineurin* (or scramble control sgRNA) without or with Flag-Optineurin phosphomutant alleles were infected with luciferase-expressing *Mtb* H37Rv (MOI 1) in four replicate wells per condition. The media were replaced daily with fresh media. (A) Monolayer luminescence was measured daily. (B) CFU were measured 3 days post-infection. The mean, SEM, and adjusted p-values from two-way ANOVA (Tukey’s multiple comparison test) are displayed. The data are representative of two independent experiments.

### Optineurin deficiency accelerates necrotic-like host cell death during *Mtb* infection

Because Optineurin deficiency enhances non-apoptotic cell death during *M. smegmatis* infection (51), we monitored macrophage viability during *Mtb* infection using real-time live-cell imaging with propidium iodide (PI) to detect membrane lysis and Caspase-3/7 CellEvent to assess apoptosis. The culture media were left undisturbed after initial infection to prevent the washing away of detached dead cells (16, 29).

At 4 days post-infection, Optineurin-deficient BMDMs exhibited a 2.1-fold increase in PI-positive cells compared to wild-type controls, demonstrating accelerated kinetic onset of necrotic-like cell death (Fig. 4A-B). By 5 days, the total numbers of PI– and CellEvent-positive cells became comparable between genotypes, indicating that while both populations eventually succumb to extensive infection-induced death, Optineurin-deficient cells possess an early kinetic vulnerability (Fig. 4A-B). This accelerated death was pathogen-dependent, as mock-infected cells maintained high viability (Fig. 4C-D). In summary, Optineurin deficiency prematurely drives macrophages toward a necrotic-like cell death pathway during *Mtb* infection (Fig. 4E).

**Fig. 4.**
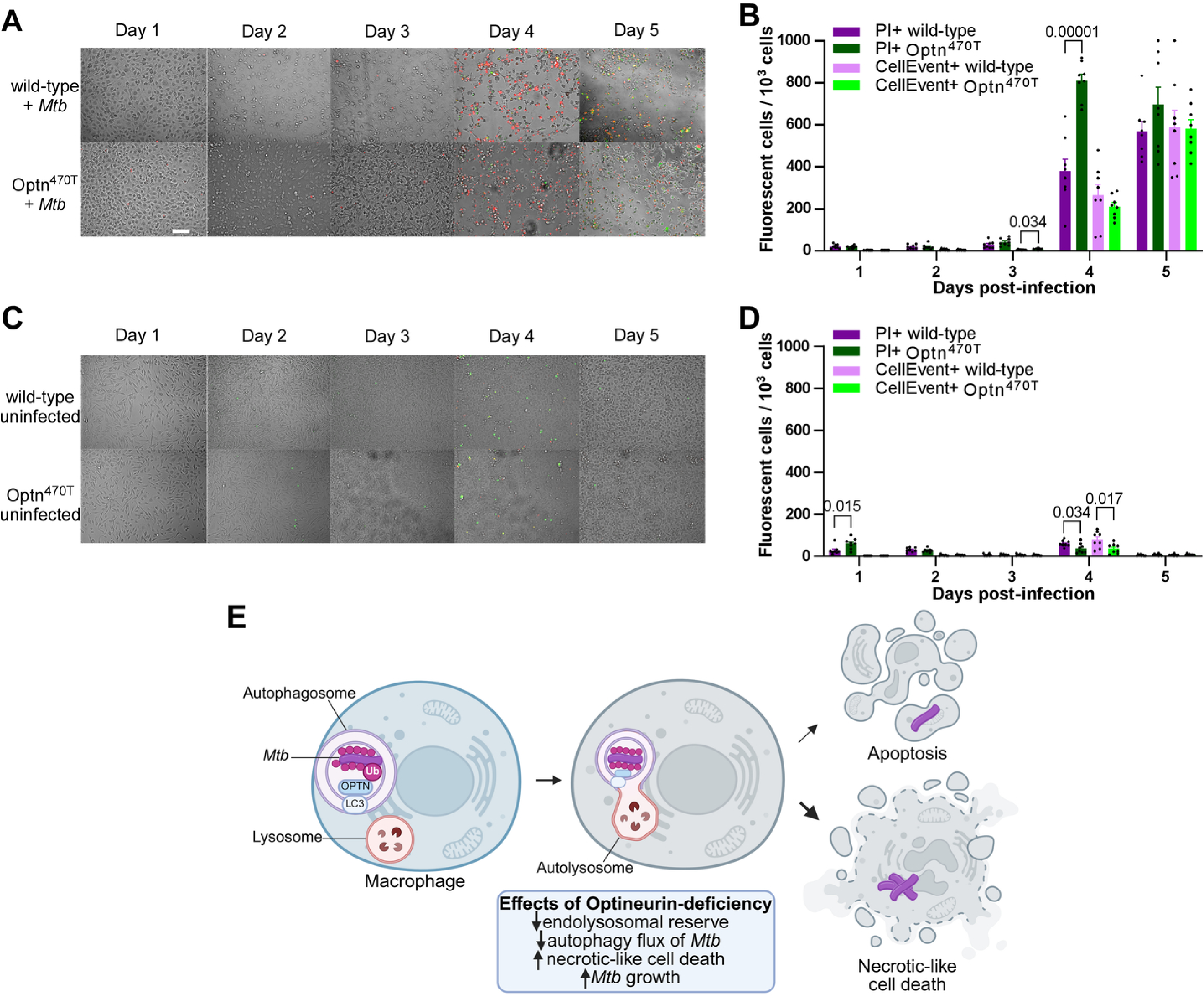
Optineurin deficiency enhances necrotic-like host cell death. BMDMs were infected with *Mtb* (MOI 1; A-B) or mock-infected (C-D) in the presence of CellEvent caspase 3/7 to detect apoptotic cells and propidium iodide (PI) for necrotic/late apoptotic cells. Fluorescence images were obtained at 20 × magnification in two positions per well in four replicate wells. The cell culture media were not replaced in these experiments to avoid disruption of dead cells in the monolayers. (A, C) Representative fluorescence and brightfield microscopy images were merged. The fluorescent cell numbers were normalized by the total number of cells in the brightfield image for each field. Mean, SEM, and statistically significant adjusted p-values from two-way ANOVA (Tukey’s multiple comparison test) comparisons are displayed. For clarity, only statistically significant adjusted p-values (p < 0.05) are shown. The white bar depicts 100 µm. The data are representative of two independent experiments. (E) Model of the effect of Optineurin deficiency on *Mtb* infection. Optineurin-deficient macrophages displayed reduced autophagy flux and endolysosomal reserve. Following *Mtb* infection, Optineurin-deficient macrophages exhibited an increase in necrotic-like host cell death and increased *Mtb* replication. Abbreviations: Ub (ubiquitin), OPTN (Optineurin). Created using BioRender.com.

### Optineurin deficiency enhances *Mtb* growth *in vivo*

To evaluate the physiological impact of Optineurin deficiency during *Mtb* pathogenesis *in vivo*, wild-type mice and homozygous Optn^470T^ mutant mice were infected with a low dose of ZsGreen-expressing *Mtb* via the aerosol route. The acute phase of infection was defined as the first 28 days post-infection, driven predominantly by innate immunity, and the chronic phase as the period beyond 28 days following adaptive immune establishment (91–93).

Lung colony-forming unit (CFU) measurements at 1-day post-infection confirmed that initial bacterial uptake was similar across genotypes (Fig. S8). Over the acute phase, Optn^470T^ mice displayed a distinct defect in bacterial restriction, exhibiting a 2.1-5.6-fold increase in lung CFU alongside a 2.3-15.8-fold increase in liver CFU and 2.6-4.2-fold increase in spleen CFU at 14, 21, and 28 days post-infection (Fig. 5A). This enhanced growth phenotype was reproducible, as a second independent experiment confirmed increased mutant organ CFU burdens at 14– and 21-day post-infection (Fig. S9).

**Fig. 5.**
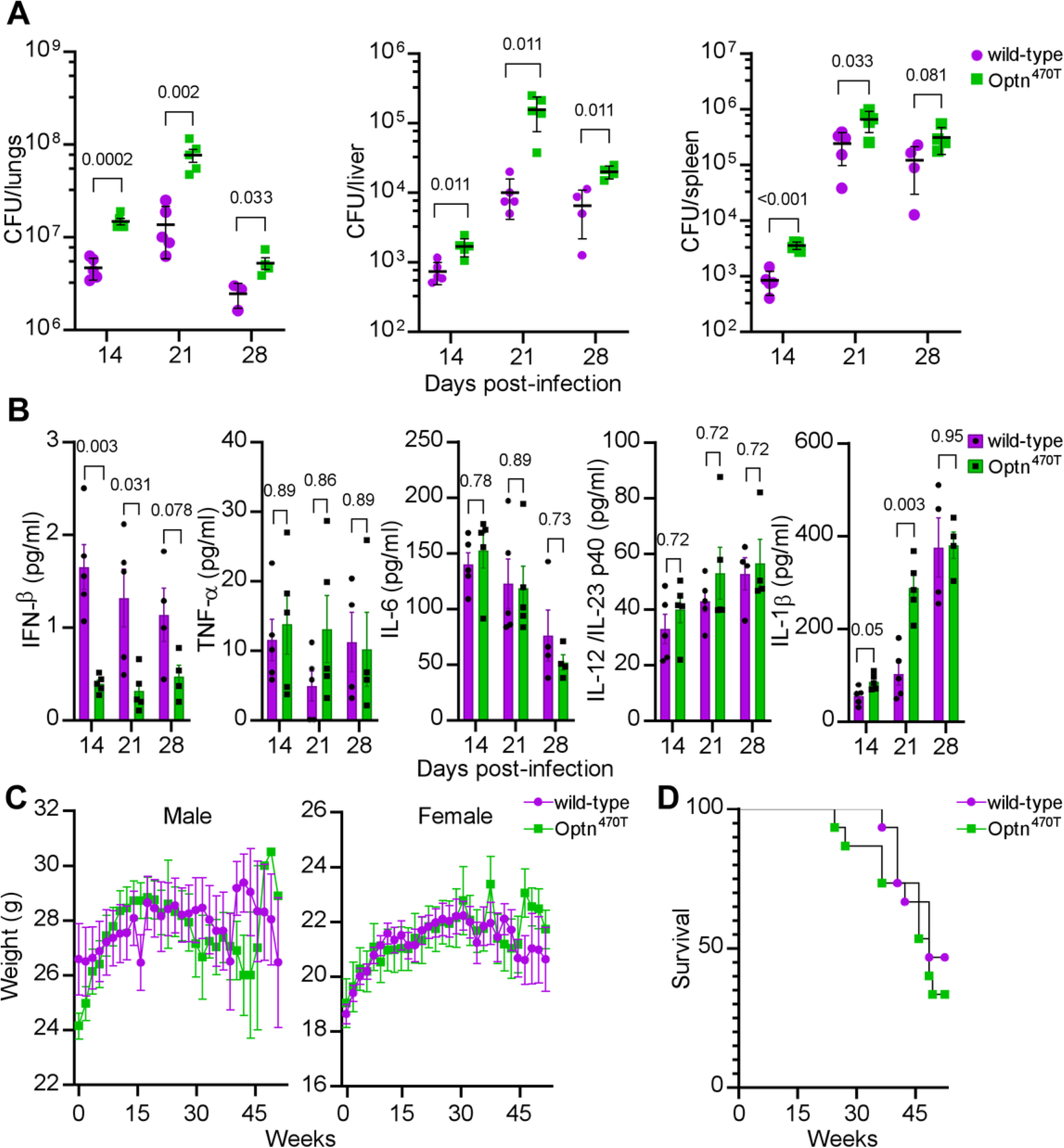
Optineurin deficiency enhances *Mtb* growth during aerosol infection. Four to five age– and sex-matched wild-type and Optn^470T^ mice were infected with ZsGreen-expressing *Mtb* by the aerosol route with 189 CFU/mouse, based on the CFU quantified from lung homogenates of mice euthanized at 1-day post-infection. (A) At 14, 21, and 28 days post-infection, CFU were quantified from the lungs, liver, and spleen. (B) Inflammatory cytokines were measured from lung homogenates. (C) The weights of the infected mice are shown. The connecting line denotes the mean weight of the mice at each time point. (D) Infected mice were monitored for death or 15% loss of maximum body weight, at which point they were euthanized. The log-rank (Mantel-Cox) test and Gehan-Breslow-Wilcoxon comparison test p-values for survival were 0.438 and 0.439, respectively. Mean (bar), SEM (error bars), and p-value from unpaired, two-tailed t-tests with Holm-Sidak multiple comparison correction are displayed.

To determine whether Optineurin deficiency altered the pulmonary inflammatory milieu, we quantified lung cytokine profiles. No differences were noted in TNF-α, IL-6, or IL-12p40 levels; however, Optn^470T^ mice displayed increased mean IL-1β at 21 days post-infection (102.6 in wild-type vs. 288.2 in Optn^470T^). Conversely, Optn^470T^ mice exhibited decreased mean IFN-β levels at 14 and 21 days (1.65 in wild-type vs. 0.39 in Optn^470T^ and 1.32 vs. 0.32, respectively; Fig. 5B). This shifted cytokine signature is consistent with the accelerated necrotic-like cell death kinetics observed in our macrophage models, which typically promotes IL-1β release while blunting interferon responses.

While prolonged Type I interferon signatures are associated with chronic active disease (94), the modest decrease in pulmonary IFN-β levels observed in Optn^470T^ mice likely reflects a cell-intrinsic signaling impairment. Because Optineurin serves as an essential structural scaffold for TBK1 activation and subsequent IRF3-mediated interferon transcription (65, 95, 96), its truncation compromises the basal host signaling axis required to mount early interferon-stimulated innate defense mechanisms (94, 97–99). Collectively, these *in vivo* findings demonstrate that loss of the C-terminal functional domains of Optineurin enhances bacterial replication and regulates the local immune microenvironment during the acute phase of *Mtb* infection.

### Optineurin deficiency does not affect survival during *Mtb* infection

Given that Optineurin deficiency accelerates acute *Mtb* growth, we monitored long-term survival and weights over 52 weeks in wild-type (n = 16) and Optn^470T^ (n = 19) mice. Log-rank analysis revealed no significant differences in long-term survival, and longitudinal animal weights were similar between genotypes (Fig. 5C-D). Thus, while Optineurin contributes to acute restriction, it is dispensable for host survival during chronic infection. This kinetic divergence likely reflects the onset of adaptive immunity, such as IFN-y-mediated macrophage activation, which remains intact in our models (Fig. 1C), acting as a dominant compensatory axis, or indicates that the moderate CFU increase falls below the threshold required to shorten survival in this low-dose aerosol model.

### Optineurin deficiency modestly alters immune cell recruitment during acute *Mtb* infection

To test whether enhanced *Mtb* growth correlated with heightened pulmonary inflammation (11, 100, 101), we quantified the lung parenchyma occupied by inflammatory lesions at 28 days post-infection. The global lesion area was unaltered by Optineurin deficiency at this time point (Fig. S10).

Next, we used analytical flow cytometry to track intracellular ZsGreen-*Mtb* distribution and quantify cellular infiltration with higher sensitivity (16). Optineurin deficiency did not alter the relative proportions of ZsGreen-positive mononuclear subset 2 (MNC2, CD11c^hi^), CD4^+^, or CD8^+^ T cells (Fig. S11A,C). Intriguingly, Optn^470T^ lungs contained an increased proportion of alveolar macrophages (AM) at 14 days (mean of 9.3 in wild-type vs. 11.7 in Optn^470T^) and mononuclear cell subset 1 (MNC1, CD11c^lo^) at 21-days post-infection (mean of 5.5 in wild-type vs. 8.7 in Optn^470T^; Fig. S11C). Optineurin deficiency also led to a non-significant upward trend in ZsGreen-positive neutrophils (PMNs), MNC1, and NK cells (Fig. S11B,D). This myeloid lineage expansion aligns with the accelerated necrosis kinetics and elevated IL-1β observed in our cellular models.

Notably, because dead *Mtb* can retain residual fluorescent signals and cells were not sorted for viable CFU enumeration, these data do not resolve whether Optineurin deficiency increased the survival of viable bacilli within specific leukocyte populations. Nevertheless, Optineurin deficiency modestly increases the proportions of AMs and inflammatory mononuclear cells during acute *Mtb* infection.

## Discussion

While Optineurin deficiency paradoxically enhances Type I interferon during viral infections (58), we found significantly reduced pulmonary IFN-β in *Mtb-*infected Optn^470T^ mice (Fig. 5B). This confirms that the regulation of Type I interferon by Optineurin is pathogen-specific. Our results align with models showing compromised LPS-induced Type I interferon responses in Optn^470T^ BMDMs (65), Optineurin-null mice (102), and TBK-1-binding-domain deficient mice (95).

Mechanistically, excessive Type I interferon worsens *Mtb* pathogenesis by inducing neutrophil extracellular traps (103), inhibiting antimicrobial IFNy signaling (104), and driving necroptosis (105). Conversely, basal autocrine Type I interferon signaling is protective during early *Mtb* infection. This baseline activation stimulates nitric oxide synthase 2 (NOS2) and prevents macrophages from shifting toward a permissive alternative activation state (98, 99).

We hypothesize that IFN-β levels in Optn^470T^ mice drop below this protective threshold, causing the observed bacterial expansion (Fig. 5A). This signaling defect works alongside disrupted xenophagy (Fig. 2). Lacking early interferon defenses and autophagic containment, mutant macrophages fail to restrict intracellular bacterial replication, ultimately triggering an accelerated, lytic necrotic-like cell death pathway (Fig. 4).

A limitation of our model is that Optn^470T^ mice exhibit a global deficiency of the C-terminal ubiquitin-binding region (65). Therefore, we cannot determine whether enhanced pathogen growth stems from hematopoietic or non-hematopoietic compartments, or from dominant effects of the truncated Optineurin protein. However, this model remains valid for mapping Optineurin’s C-terminal, ubiquitin-dependent functions, because CRISPR/Cas9-engineered Optineurin-knockout CIM lines (Fig. 1B) recapitulate the accelerated *Mtb* replication seen in Optn^470T^ BMDMs (Fig. 1C).

Impaired *Mtb* control in macrophages expressing phosphodeficient *Optineurin* alleles confirms that post-translational signaling regulates host defense (106). Among the three growth-enhancing phosphomutants, human S513 site (murine S530) inside the UBAN domain enhances ubiquitin affinity (106), while human S177 (murine S187) within the LIR motif optimizes LC3 binding to drive cargo clearance (40, 44, 60, 107). In contrast to these phosphorylation sites with established functions, the third critical site in the zinc finger domain (human S539, murine S556) has no known canonical function. Because of its proximity to the UBAN domain, S556 phosphorylation may regulate secondary ubiquitin binding or receptor oligomerization, which requires structural validation (108).

Our study highlights the nonredundant roles of selective autophagy receptors during *Mtb* infection. Unlike Tax1bp1 deficiency, which limits *Mtb* replication (16), or p62 deficiency, which has negligible effects (33), Optineurin deficiency accelerates *Mtb* growth, arrests autophagy flux, and triggers necrotic-like host cell death. These differences likely stem from structural variations in their ubiquitin-binding domains (108, 109). Optineurin prefers linear ubiquitin, a regulator of inflammasome assembly (110), whereas Tax1bp1 binds K63– and K48-linked ubiquitin chains to drive NF-κB inflammatory signaling (109, 111). Consistently, Tax1bp1 deficiency blunts NF-κB inflammation (16). Furthermore, Tax1bp1 and Optineurin assemble distinct signalosomes. Tax1bp1 interacts with A20 (112), Itch (113), and RNF11 (114), while Optineurin binds CYLD (115) and NEMO/IKKy (116), transmitting nonredundant signals.

Although *Mtb* triggers mitochondrial damage and ER stress, Optineurin’s roles in mitophagy or ER homeostasis are unlikely drivers of the growth phenotype we report here. *Mtb* exploits mitophagy to enhance replication (117). Thus, mitophagy disruption should restrict rather than accelerate bacterial growth. Furthermore, while ER stress traditionally triggers apoptosis (118), Optn^470T^ BMDMs showed no apoptotic expansion (Fig. 4A-B). Therefore, these collateral pathways are unlikely to account for the accelerated *Mtb* growth observed in our study.

In summary, Optineurin is a major regulator of host defense against *Mtb*, and its deficiency disrupts xenophagic flux and compromises bacterial restriction *in vitro* and *in vivo.* Therapies that augment Optineurin activation or mimic its phosphorylation states represent promising host-directed strategies for restoring autophagic clearance of intracellular *Mtb*.

## Material and methods

### Ethics statement

Animal infections were performed in accordance with the animal use protocol (AN192778) approved by the Institutional Animal Care and Use Program at the University of California, San Francisco, in compliance with federal regulations established by the National Research Council and the National Institutes of Health.

### Mice and macrophage cell lines

Wild-type C57BL/6J mice were purchased from Jackson Laboratories. Optn^470T^ mice were obtained from Ivana Munitic, University of Rijeka, Croatia (65). Optineurin-deficient mice were rederived upon transfer to UC San Francisco. Both wild-type and Optn^470T^ primary murine bone-marrow-derived macrophages (BMDMs; Table 1) were prepared by flushing the femurs from 8– to 15-week-old mice. Extracted bone marrow cells were differentiated for 7 days and cultured in DMEM (Dulbecco’s Modified Eagle Medium), high glucose supplemented with 20% FBS (fetal bovine serum), 2 mM glutamine, 0.11 mg/ml sodium pyruvate, and 15% MCSF (macrophage colony-stimulating factor) derived from 3T3-MCSF cells (BMDM media). Progenitor CIMs (Cas9+ conditionally-immortalized macrophages) were maintained in suspension in non-treated tissue culture-treated flasks with RPMI (Roswell Park Memorial Institute) media supplemented with 10% FBS, 2% GM-CSF (granulocyte-macrophage colony-stimulating factor) supernatant produced by a B16 murine melanoma cell line, 2 mM L-glutamine, 1 mM sodium pyruvate, 10 mM HEPES, 43 uM β-mercaptoethanol, and 2 µM β-estradiol before differentiation. CIMs were washed with PBS (phosphate-buffered saline) + 1% FBS twice to remove β-estradiol, resuspended in macrophage media, and seeded with 1.56 × 10^6^ cells in a non-treated 10 cm plate to differentiate. After 3 days, 4 ml of macrophage media was added, and on day 7, harvested with 0.05% trypsin for assays.

**Table 1.**
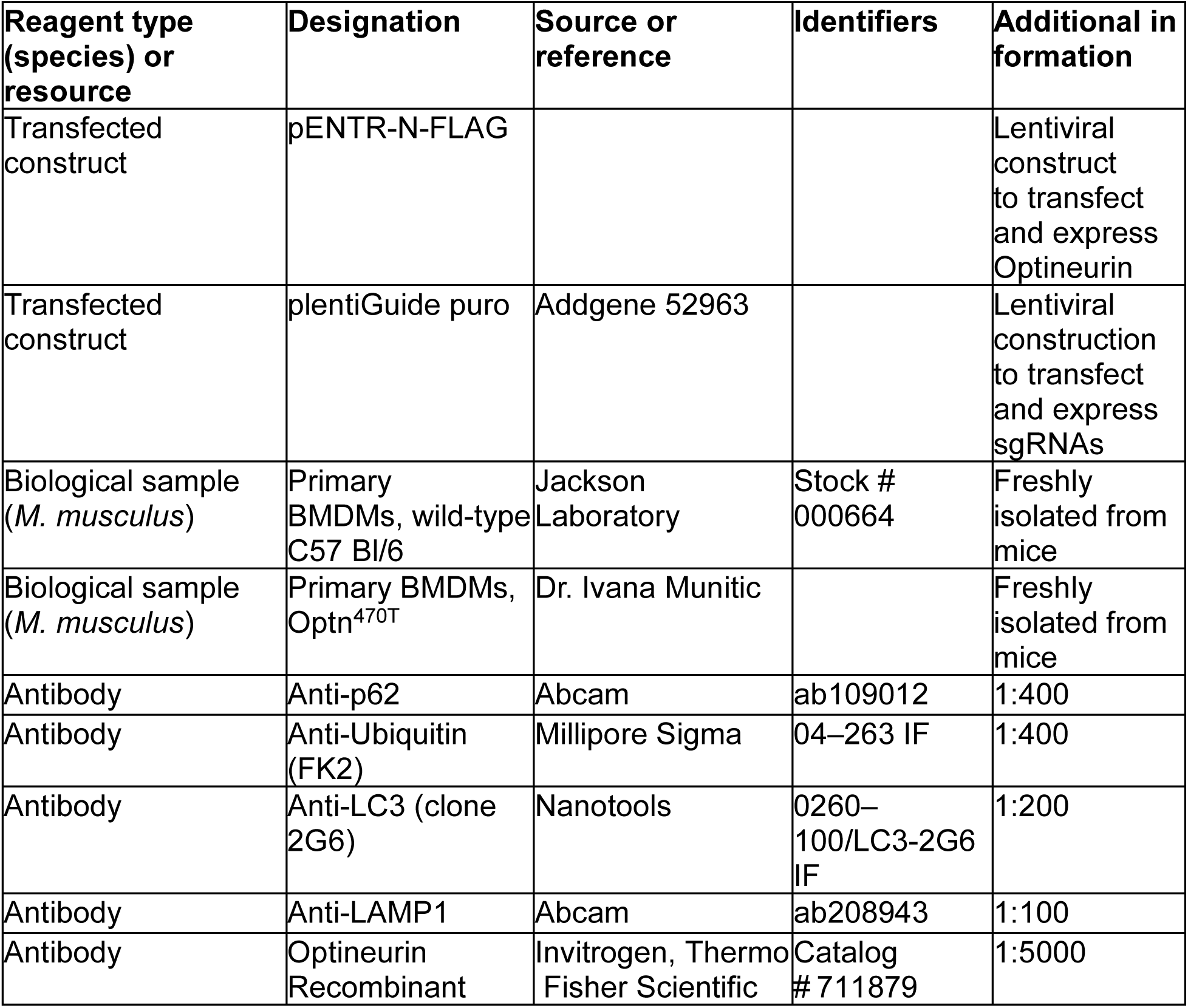
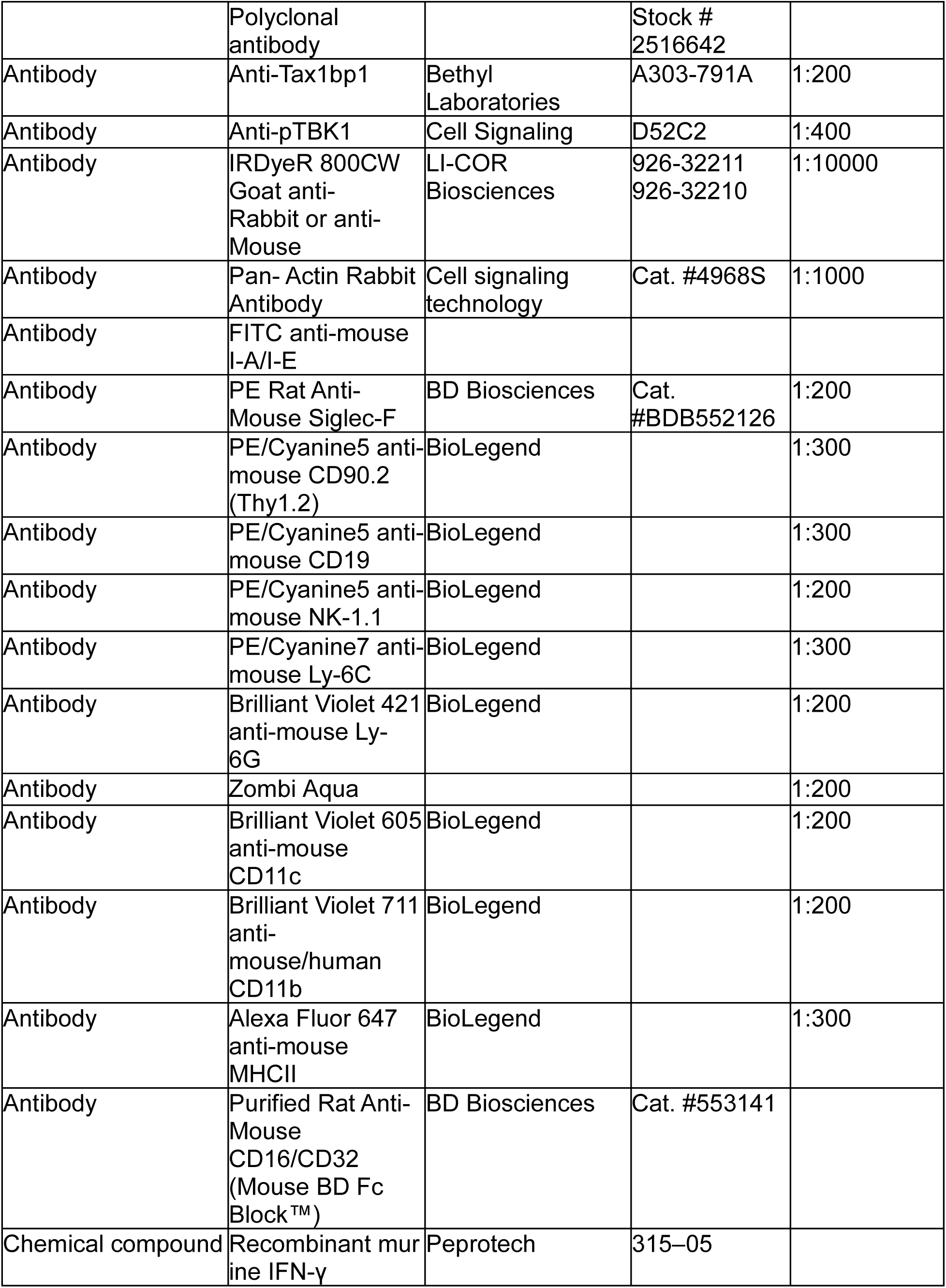
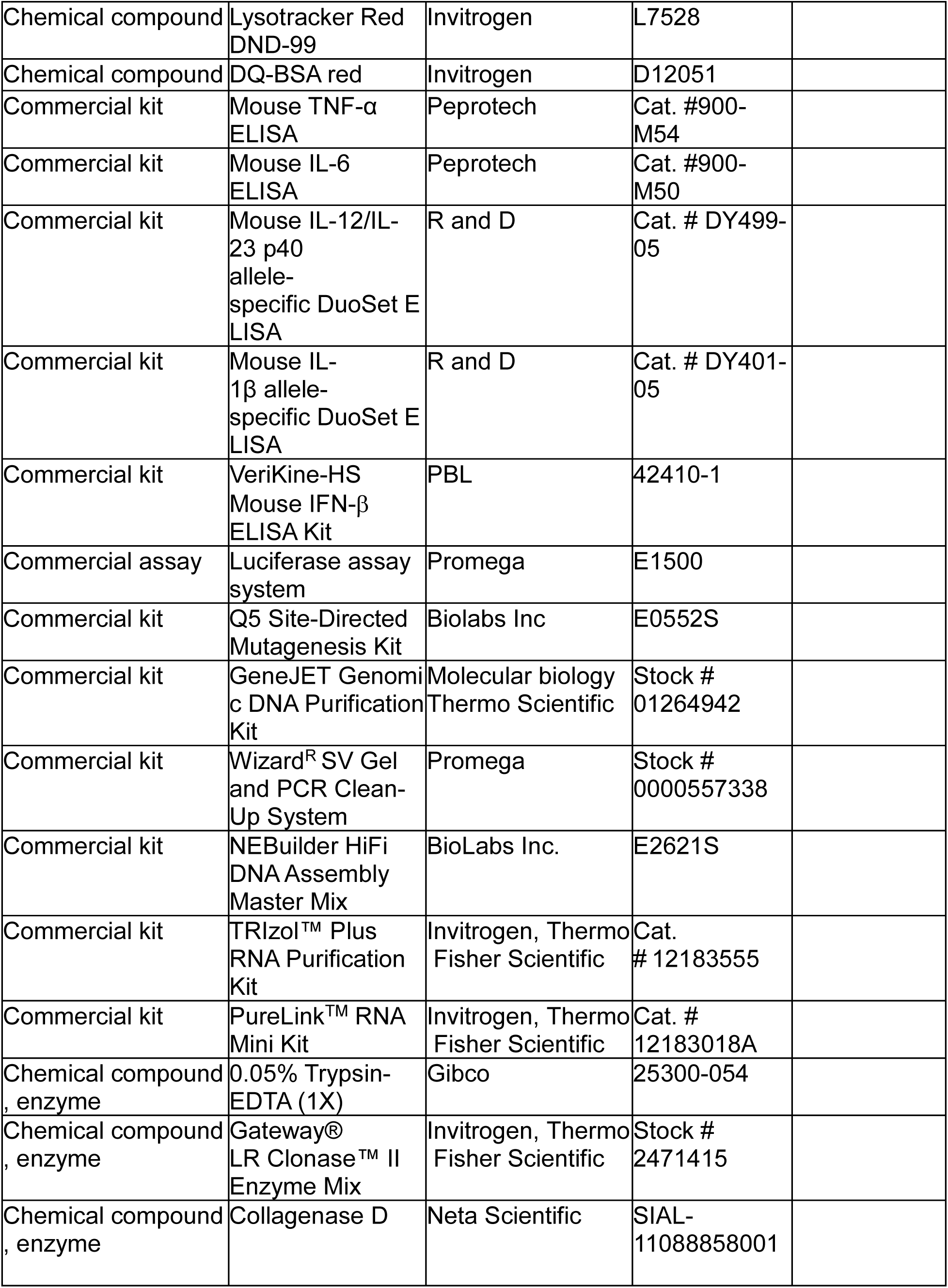
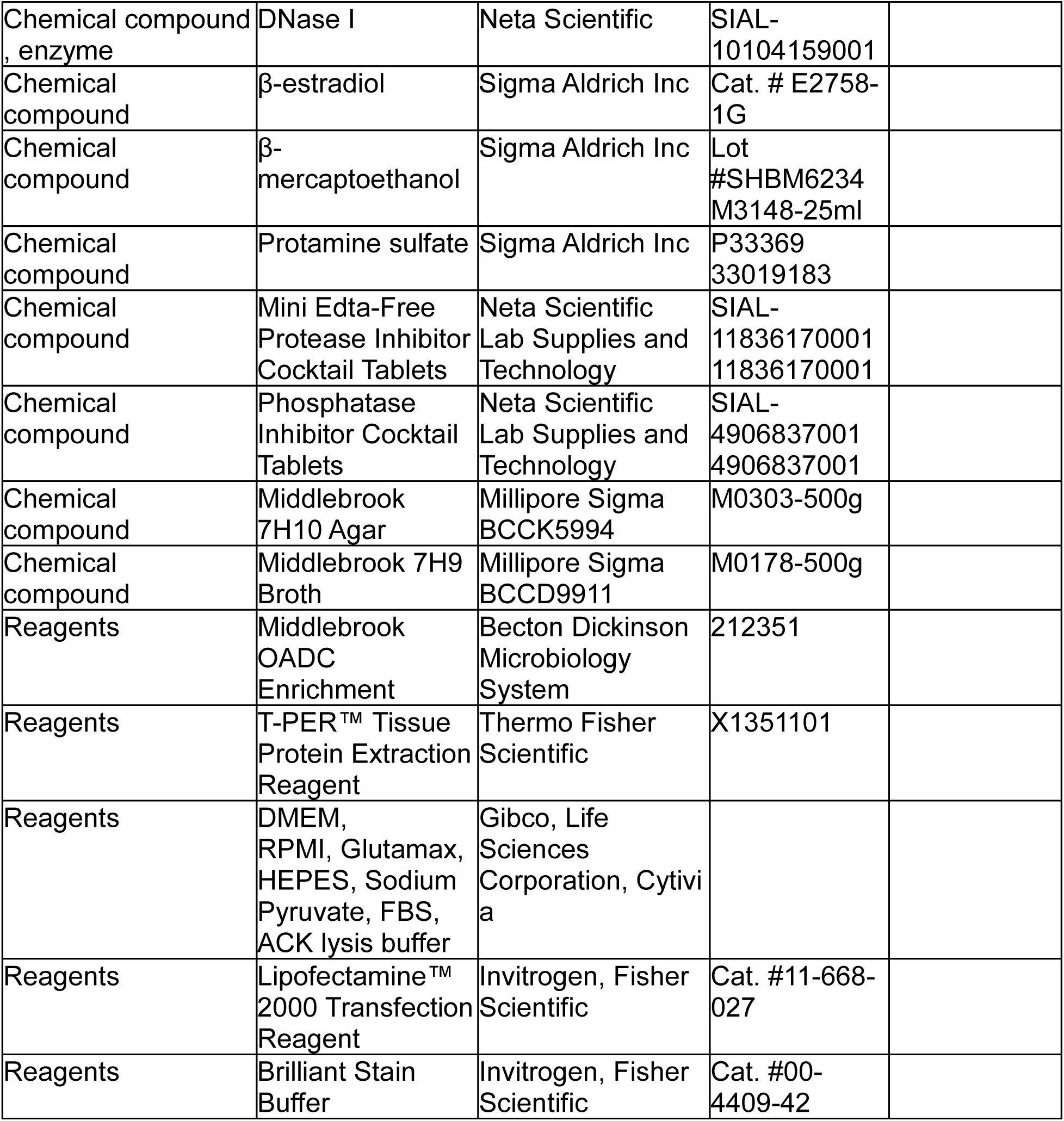
Reagents used in this study.

### Bacterial strains

*M. tuberculosis*: Wild-type Erdman, H37Rv expressing the luxCDABE operon (pMV306hsp-LuxG13, Addgene #26161), and Erdman (pMV261::ZsGreen) were used for infection. *M. tuberculosis* was cultured to mid-log phase in 7H9 broth media supplemented with 10% Middlebrook OADC, 0.5% glycerol, and 0.05% Tween80 at 37 °C with orbital shaking at 100 rpm.

### Lentiviral transduction of CRISPR sgRNAs to generate *Optineurin* knockout progenitor conditionally immortalized macrophages (CIMs)

Single oligonucleotide guides (CRISPR sgRNAs) for scramble control and Optineurin were designed from the Brie library and cloned into pLenti-gRNA hygro (Addgene plasmid #104991) using the primers P1/P2 (Scramble control; Table 2) and P3/P4 (Optineurin exon 2). LentiX 293T cells (1.4×10^6^ cells/well in TC-treated 6-well plates) were cotransfected with vectors encoding scramble control or Optineurin guides. Cas9-expressing CIM progenitors (5.0 × 10^5^ cells/well in a 6-well plate) were transduced with lentiviral particles at 1000 × g for 2 h at 32 °C with protamine sulfate (10 µg/ml). After two days of transduction, cells were treated with hygromycin (12 µg/ml) for 4 days.

**Table 2.**
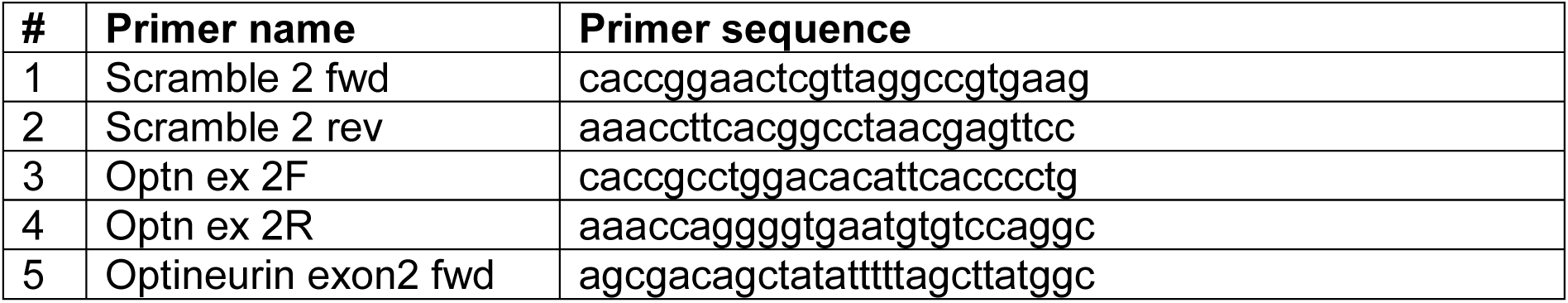
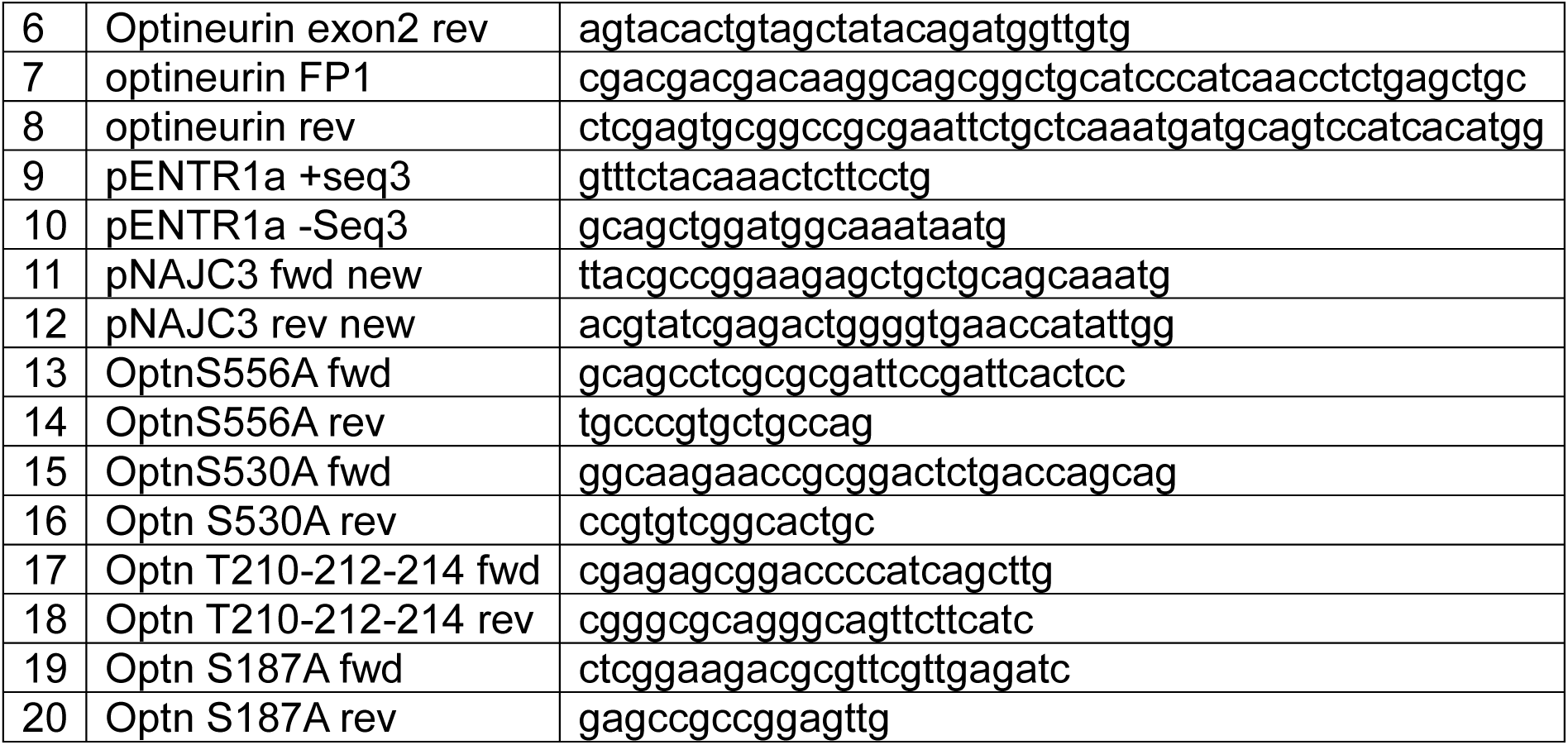
Primers used in this study.

### Isolation of a monoclonal cell population by limiting dilution

In a 96-well plate, 1.6 × 10^4^ polyclonal edited cells were serially diluted in 50 µl of progenitor media and incubated at 37 °C with 5% CO_2_ for 24 h. The plates were examined under a microscope to identify wells containing a single cell. Every 5 days, each well received 50 µl of progenitor medium, with β-estradiol concentrations maintained at 2 µM. After 20 days, genomic DNA was extracted using the GeneJET Genomic DNA purification kit. Genomic sites encompassing targeted regions were amplified by PCR (polymerase chain reaction) using Q5 high-fidelity DNA polymerase (NEB) using primers P5/P6 (Optineurin exon 2). The resulting PCR product was purified using the Wizard SV Gel and PCR clean-up system and sequenced with primer P5. Inference CRISPR Edits (ICE) from Sanger sequencing trace data (Synthego) were used to determine population-level genome editing.

### Cloning Optineurin alleles for complementation and phosphomutant analysis RNA extraction from BMDM macrophages

Total RNA was isolated using the PureLink^TM^ RNA Mini Kit, and DNA was removed using on-column PureLink DNAse, as per the manufacturer’s protocol.

### Synthesis of first-strand cDNA

First, 5 µg of total RNA, 1 µl of oligo(dT) primers for the eukaryotic template, 1 µl of 10 mM dNTP mixture, and DEPC-treated water were combined in a 0.5 ml microcentrifuge tube. The reaction mixture was then incubated at 65 °C for 5 min and placed on ice for 1 minute. The cDNA synthesis mixture was prepared by mixing 2 µl of 10X RT buffer, 4 µl of 25 mM MgCl_2_, 2 µl of 0.1M dNTPs, 1 µl of RNase OUT, and 1 µl of Superscript III RT solution. cDNA synthesis mixture (10 µl) was added to each RNA/primer mixture and collected by centrifugation. The samples were incubated for 50 min at 50 °C. The reaction was terminated at 85 °C for 5 min and chilled on ice. RNase H was added to each tube, and the samples were incubated for 20 min at 37 °C.

### Cloning of FLAG-tagged Optineurin alleles into entry and destination vectors

PCR products were generated for *Optineurin* by amplifying the Optineurin coding region from the cDNA using the primer pair P7/P8. PCR products were cloned into pENTR-N-FLAG, encoding a 3x N-terminal FLAG tag (15), using Fsp-I and ligation-independent cloning (NEBuilder HiFi DNA Assembly Reaction Protocol). The plasmids were isolated and sequenced using the primers P9/P10. The Q5^®^ Site-Directed Mutagenesis Kit was used to engineer synonymous mutations at the sgRNA target site of *Optineurin* exon 2 using primers (P11/P12), thereby avoiding target recognition by the sgRNA in complemented strains. Site-directed mutagenesis of Optineurin phosphosites was performed using the following primer pairs: P13/P14 (S556A), P15/P16 (S530A), P17/P18 (T210A, T121A, T214A), and P19/P20 (S187A). Gateway LR reaction was used to clone Optineurin receptor mutants into the pLenti-CMV Puro DEST (w118-1) lentiviral Gateway destination vector.

### Transfection and lentiviral transduction of FLAG-tagged Optineurin alleles in *Optineurin-*deficient CIMs

As described for the generation of the Optineurin-edited polyclonal mutants, lentivirus was generated for the transduction of monoclonal Cas9-expressing *Optineurin*^-/-^ (exon 2) CIM progenitors with vectors expressing FLAG-tagged Optineurin alleles. After two days of transduction, cells were selected in puromycin (12 µg/ml) for 4 days. Puromycin-resistant cells were expanded and maintained for further use.

### Western blots

30 µg of protein lysate was subjected to SDS-PAGE (BioRad Miniprotean TGX 4–20%) and transferred onto a nitrocellulose membrane. The membrane was blocked with Odyssey blocking buffer and incubated with anti-Optineurin antibody at a dilution of 1:5000, at 4 °C overnight. After the membranes were probed with a secondary antibody (goat anti-rabbit 1:10000) for 1 h, the membrane was imaged on an Odyssey scanner (Li-Cor). The same nitrocellulose membrane was re-probed with α-actin mouse monoclonal antibody. LC3 immunoblot was performed as previously described (16).

### *Mtb* growth assays

Differentiated CIMs or BMDMs were seeded at 60,000 cells per well onto white, clear-bottom Cell Bind 96-well plates (Corning) in macrophage media 24 h before infection. Mid-log *M. tuberculosis* culture was pelleted at 3,500 rpm and washed twice with PBS to remove excess Tween 80, gently sonicated to disperse clumps, and resuspended in phagocytosis infection media (DMEM supplemented with 5% horse serum albumin). Bacterial density was evaluated by measuring absorbance at 600 nm, and the multiplicity of infection (MOI) was adjusted to macrophage cell density. For *Mtb* growth assays, macrophages were infected at an MOI of 1 by removing the media from the cells and adding bacterial suspensions in phagocytosis media, and then “spinfected” at 1,000 rpm for 10 min. Afterward, the infection media was removed, and fresh macrophage media was added. The bacterial luminescence signal was measured at the time of infection and every day thereafter (4 days post-infection) following media changes. In experiments including IFN-y, the media were replaced daily with fresh media containing 7.5 ng/ml of murine IFN-y. Bacterial growth measurements were normalized to luminescence readings at day 0 for each well and represented as fold changes in luminescence compared to day 0. *Mtb* CFU enumeration experiments were performed by lysing macrophages with 0.1% Triton X-100 solution on day 0 and day 4, and serial dilutions were performed in PBS with 0.05% Tween-80. Diluted samples were plated on Middlebrook 7H10 agar. CFUs were counted after incubation of plates at 37 °C for 3 weeks.

### Cytokine measurements

Homogenized mouse lung tissue (700 µl) was mixed with an equivalent volume of T-PER lysis buffer solution (with Protease Inhibitor Cocktail Tablet and Phosphatase Inhibitor Cocktail Tablet), vortexed, and incubated at 4 °C for 20 min. The samples were centrifuged at 12000 × g for 10 min and filtered through a 0.22 µm filter. Cytokines were measured in the supernatants according to the manufacturers’ instructions for the ELISA (Enzyme-Linked Immunosorbent Assay) kits.

### Mycobacterium tuberculosis aerosol infection

Mice were infected with *Mtb* (pMV261:: ZsGreen) via the aerosol route, using an inhalation exposure system unit (Glas-Col). Our target infectious dose was ∼100 CFU/mouse, quantified on day 1 post-infection by plating whole-lung homogenates from 3 to 5 mice onto Middlebrook 7H10 agar. To determine the bacterial load throughout the infection, lungs, spleen, and liver were harvested on day 14, 21, and 28, homogenized, and serial dilutions were plated on Middlebrook 7H10 agar. CFUs were counted after incubation of plates at 37 °C for 3 weeks.

### Flow cytometric analysis

Mouse lungs were harvested and processed into single-cell suspensions and stained with the antibody panels for immune cells as described previously (16). For flow cytometry, samples were analyzed using a BD LSRII. For immunophenotyping and flow cytometry analysis, statistical differences between wild-type and Optn^470T^ mouse lungs were assessed for each cell subset using planned, independent, two-tailed unpaired t-tests. Because each immune cell lineage represents an independent biological recruitment mechanism during infection, p-values were evaluated without multi-variable global adjustments to preserve sensitivity for discovery-driven leukocyte profiling. A p-value < 0.05 was considered statistically significant.

### Live cell imaging

BMDMs were infected with wild-type Erdman *Mtb* at an MOI of 1. To measure necrosis and apoptosis, 1 µg/ml of propidium iodide (Life Technologies) and two drops per milliliter of CellEvent Caspase-3/7 Green ReadyProbes reagent (Invitrogen) were added to the media at the beginning of the infection. In these experiments, the cell culture media were changed immediately after infection, but not on subsequent days, to avoid disruption of the cell culture monolayer (16, 62). Fluorescence and phase contrast images were obtained at 20 × magnification using a Keyence BZ-X 700 microscope. Images were obtained daily in four technical replicates per condition and at two positions per well. Quantification of necrotic and apoptotic cells was performed using ImageJ. Cell counts in brightfield images were quantified using a Python script that counted cells based on specified distances and cell diameters.

### Immunofluorescence microscopy

Wild-type and Optn^470T^ BMDMs were seeded at a density of 50,000 cells/well in 96-well plates in quadruplicate wells and incubated overnight at 37 °C with 5% CO_2_. Macrophages were infected with *Mtb* at an MOI of 2 and incubated at 37 °C with 5% CO_2_. At 8– or 24-h post-infection, the infected macrophages were fixed with 4% PFA, the monolayers were washed with PBS, and the samples were stored at 4 °C. For experiments including bafilomycin at final concentrations of 50-1000 nM, the chemical was added at 21 h post-infection. For LC3 immunostaining, samples were additionally fixed with ice-cold methanol for 10 min. Permeabilization and blocking were performed in PBS with 2% BSA and 0.1% saponin or 0.3% Triton X-100 (for LC3). Cells were stained with primary antibodies at room temperature for 1.5 h or 4 h (for LC3) and secondary antibodies at 1:4000 or 1:1000 (for LC3). Cells were washed three times for 5 min in PBS and stained with DAPI (4’,6-diamidino-2-phenylindole; 1:1000) and anti-mouse or –rabbit Alexa Fluor 647 (Invitrogen) for 1 h. Cells were washed three times for 5 min in PBS. In experiments including Lysotracker, macrophages were treated with 100 nM Lysotracker at 5 h post-infection. Three h later, the monolayers were chemically fixed, washed with PBS, and immediately imaged. In experiments including DQ-BSA, the macrophages were treated with bafilomycin at a final concentration of 300 nM at 5 h post-infection, DQ-BSA (10 µg/ml) at 7 h post-infection, followed by fixation at 8 h post-infection.

### Automated confocal microscopy and image analysis

Imaging was performed using an Opera Phenix High-Content Screening System confocal microscope (PerkinElmer) with a 63 × magnification lens. Colocalization analysis was performed using PerkinElmer Harmony software. Maximum intensity image projections from multiple z-stacks were generated as previously described (16). Bacterial and autophagy markers were quantified using method C (for Tax1bp1, polyubiquitin, LC3, and phosphorylated-TBK1) or method A (for p62) to minimize the selection of non-specific background staining. Bacteria with at least one pixel of overlap with autophagy marker spots were counted as colocalization events. Percent colocalization was defined as the number of colocalization events divided by the total number of bacteria. Percent colocalization, corrected spot intensity, and puncta per cell were calculated for each well from all the images obtained in the well using the evaluation module.

### RNA purification for differential gene expression analysis

Three independent biological replicate infection experiments were performed for RNA isolation using BMDMs harvested on three separate occasions from wild-type and Optn^470T^ mice. BMDMs were infected with wild-type *Mtb* (MOI 2) or mock-infected by “spinfection” as described under *Mtb* growth assays in 96-well plates. At 24 h post-infection, the monolayers from 5-10 wells per experimental condition were washed with PBS, the macrophages were lysed with 100 µl of Trizol reagent per well, and the samples from each condition were pooled together for RNA purification using the Direct-zol RNA miniprep kit (Zymo Research) following the manufacturer’s protocol. RNA was eluted in IDTE buffer. Purified RNA was analyzed using a RNA bioanalyzer for RNA Integrity Number (RIN) measurement, stabilized with SEQguard Dino Preserve, and shipped to Plasmidsaurus at ambient temperature. For sequencing library preparation, mRNA was converted into complementary DNA (cDNA) via reverse transcription and second-strand synthesis, followed by tagmentation, library indexing, and amplification. Differential gene expression was captured using 3’ end counting.

### Differential gene expression analysis

Quality of the fastq files was assessed using FastQC v0.12.1. Reads were then quality filtered using fastp v0.24.0 with poly-X tail trimming, 3’ quality-based tail trimming, a minimum Phred quality score of 15, and a minimum length requirement of 50 bp. Quality-filtered reads were aligned to the reference genome using STAR aligner v2.7.11 with non-canonical splice junction removal and output of unmapped reads, followed by coordinate sorting using samtools v1.22.1. PCR and optical duplicates were removed using UMI-based deduplication with UMIcollapse v1.1.0. Alignment quality metrics, strand specificity, and read distribution across genomic features were assessed using RSeQC v5.0.4 and Qualimap v2.3, with results aggregated into a comprehensive quality control report using MultiQC v1.32. Gene-level expression quantification was performed using featureCounts (subread package v2.1.1) with strand-specific counting, multi-mapping read fractional assignment, exons and three prime UTR as the feature identifiers, and grouped by gene_id. Final gene counts were annotated with gene biotype and other metadata extracted from the reference GTF file. Sample-sample correlations for sample-sample heatmap and PCA were calculated on normalized counts (TMM, trimmed mean of M-values) using Pearson correlation. Differential expression was performed using edgeR v4.0.16 using standard practice including filtering for low-expressed genes with edgeR::filterByExprwith default values.

### Data access statement

RNAseq metadata is accessible in the Gene Expression Omnibus (GEO) Accession # GSE344459.

## Acknowledgements

We thank Dr. Ady Steinbach for helpful insights regarding microscopy data analysis, Dr. Cherilyn Elwell for critical review of the manuscript, and Dr. Jeffrey Cox for valuable conceptual insights regarding Optineurin. The research reported in this publication was supported by the National Institute of Allergy and Infectious Diseases of the National Institutes of Health under Award Number R01AI194696. The content is solely the responsibility of the authors and does not necessarily represent the official views of the National Institutes of Health. Experimental data were produced using an Opera Phenix microscope in the Parnassus Center for Advanced Technology at UCSF.

